# The way of the light: how visual information reaches the auditory cortex in congenitally deaf adults

**DOI:** 10.1101/2022.01.25.477765

**Authors:** Giuliana Martinatti Giorjiani, Zohar Tal, Hamed Nili, BI Yanchao, Fang Fang, Jorge Almeida

**Affiliations:** Proaction Laboratory, Faculty of Psychology and Educational Sciences, University of Coimbra. Portugal; Center for Research in Neuropsychology and Cognitive Behavioural Intervention (CINEICC), Faculty of Psychology and Educational Sciences, University of Coimbra, Portugal; Wellcome Centre for Integrative Neuroimaging, University of Oxford, United Kingdom; Department of Excellence for Neural Information Processing, Center for Molecular Neurobiology (ZMNH), University Medical Center Hamburg-Eppendorf (UKE), 20251, Hamburg Germany; State Key Laboratory of Cognitive Neuroscience and Learning and IDG/McGovern Institute for Brain Research, Beijing Normal University, Beijing, China; Beijing Key Laboratory of Brain Imaging and Connectomics, Beijing Normal University, Beijing, China; Chinese Institute for Brain Research, Beijing, China; School of Psychological and Cognitive Sciences and Beijing Key Laboratory of Behavior and Mental Health, Peking University, Beijing, China; IDG/McGovern Institute for Brain Research, Peking University, 100087 Beijing, China; Peking-Tsinghua Center for Life Sciences, Peking University, 100087 Beijing, China

**Keywords:** Congenital deafness, Neuroplasticity, Superior Colliculus, Inferior Colliculus, fMRI, Dynamic Causal Modeling, Representational Connectivity Analysis

## Abstract

Human and animal studies on cross-modal plasticity under congenital deafness suggest that early auditory cortex plays a significant role in the processing of visual information when congenitally deprived from its typical (auditory) input. However, the pathway by which early auditory cortex is fed with visual information is still understudied. Here we focused on addressing how visual information reaches the auditory cortex under congenital deafness. We put forth a mechanistic model that proposes that different corticocortical and subcortical connections play a central role in rerouting visual information to the early auditory cortex of congenitally deaf individuals. Specifically, we show, using Representational Connectivity Analysis (RCA) and Dynamic Causal Modeling (DCM), that connections from the right superior colliculus to the right inferior colliculus, as well as connections from right early visual cortical regions to the right early auditory cortex play a role in rerouting visual information to early auditory cortex in congenitally deaf individuals. These findings shed light on how visual information reaches the early auditory cortex of deaf individuals - specifically, they suggest that neuroplasticity reshapes subcortical connections in order to re-route visual information to the auditory stream.

## Introduction

The effects of congenitally sensorial deprivation in the brain have been widely investigated in both human and non-human populations. Different studies have consistently shown that the brain undergoes several changes that include the recruitment of the sensorially-deprived areas for the processing of sensory stimuli from other modalities (e.g., Collignon et al., 2009; Lomber et al., 2010; Merabet & Pascual-Leone, 2010; Bottari et al., 2014; Almeida et al 2015; Hribar et al., 2020; for review see Bell et al., 2019). Congenital sensorial deprivation has been also found to affect behavioral performance on tasks that depend on the non-affected sensory inputs (e.g., Hong & Song, 1991; Levänen & Hamdorf, 2001; Voss et al. 2015; Almeida et al., 2018). One aspect that figures critically in our understanding of this type of long-term neuroplasticity is how non-affected sensorial information is re-routed to sensorially deprived areas. Here we will address this question, and focus on the route taken by visual information to the auditory cortex of congenitally deaf humans.

Congenital deafness has been associated with changes at the behavioral and neural levels (e.g., Neville & Lawson, 1987; Loke & Song, 1991; Stivalet et al., 1998; Lomber et al., 2010; Shiell et al., 2014; Almeida et al., 2015; 2018; Simon et al., 2020; for reviews see Amaral & Almeida, 2015; Bell et al., 2019). Specifically, numerous studies have systematically shown that early deafness leads to enhanced performance over different visual features including motion detection (e.g., Bosworth and Dobkins, 2002; Shiell et al., 2014; Hauthal et al., 2013; Simon et al., 2020), and the processing of stimuli in central (Stivalet et al., 1998), and peripheral visual fields (e.g., Neville & Lawson, 1987b; Lore & Song, 1991; Nava et al., 2008; Dye et al., 2009; Codina et al., 2017; Almeida et al., 2018). For example, Lore and Song (1991) showed that congenitally deaf individuals, when compared to hearing individuals, were faster at detecting visual stimuli presented in the periphery of the visual field, and that attention-related ERPs to peripheral stimuli were much larger in deaf than in hearing individuals (Neville & Lawson, 1987b). Almeida and colleagues (2018) further characterized these behavioral compensatory processes and tested whether motion detection was modulated as a function of visual field localization. Specifically, using a direction of motion discrimination task, they found that the performance of deaf individuals was better for stimuli that were presented in the periphery within the horizontal plane, when compared to those presented in the vertical plane or the central visual field. This visual field asymmetry was observed only in the congenitally deaf group, while hearing controls performed better for centrally presented stimuli.

Concomitantly, there are neural changes due to congenital deafness. One obvious candidate, and one area that has been singled out in both human and non-human studies, is the auditory cortex (e.g., Lomber et al., 2010; Vachon et al., 2013; Bottari et al., 2014; Scott et al., 2014; Almeida et al., 2015; Shiell et al., 2015; Bola et al., 2017; Seymour et al., 2017; Simon et al., 2020; for a review see Alencar et al., 2019; Bell et al., 2019; Amaral & Almeida, 2015). Electrophysiological studies across numerous species provide direct evidence for cross-modal plasticity at the single unit level. Specifically, these studies have found an increase in the number of neurons in the auditory cortex that respond to visual and somatosensory stimuli (Hunt et al., 2006; Meredith & Lomber, 2011; Meredith et al., 2012; Land et al., 2016). Furthermore, Lomber and Colleagues (2010) showed that enhanced performance on visual motion detection and localization tasks exhibited by deaf cats, when compared to hearing cats, is causally related to processing within the auditory cortex. A reversible deactivation of different parts of the auditory cortex has further shown that different visual tasks can be specifically localized at different portions of the deprived auditory cortex (e.g., motion detection and visual localization; Lomber et al., 2010).

Regarding deaf humans, neuroimaging studies have shown recruitment of auditory cortical areas in response to various visual stimuli (Finney, et al., 2001;2003; Fine et al., 2005; Vachon et al., 2013; Scott et al., 2014; Almeida et al., 2015; Bola et al., 2017). Visually evoked activity in deaf was found in the early auditory cortex (including BA 41 and 42; Finney et al., 2001; Fine et al., 2005; Vachon et al., 2013; Almeida et al., 2015) as well as in associative and higher auditory areas (Sadato et al., 2004; Cardin et al., 2013; Scot et al., 2014; Shiell et al., 2015 Benetti et al., 2017; Bola et al., 2017). Similar to the findings from animal studies, human cross-modal neural plasticity reflects the behavioral enhancement of different visual features (Bottari et al., 2011; Almeida et al., 2015; Seymour et al., 2017). For example, Almeida and colleagues (2015) showed that the early auditory cortex of deaf, but not hearing individuals, encodes visual information presented in visual stimuli locations within the horizontal meridian, in line with the behavioral advantage presented by deaf individuals in a motion detection task (Almeida et al, 2018).

These results pose a question as to how visual information reaches the auditory cortex of congenitally deaf individuals. That is, how is visual information re-routed towards the auditory cortex of congenitally deaf individuals? The range of the possibilities is certainly limited by the typical route of visual information to the visual cortex: visual information may reach the auditory cortex either through corticocortical connections from early visual cortex, or through subcortical connections with visual subcortical areas (e.g., Superior Colliculus and Lateral Geniculate Nucleus). These connections may relay visual information directly to the deprived auditory cortex, or indirectly through typical auditory relay stations (e.g., Inferior Colliculus and Medial Geniculate Nucleus).

Extant human and non-human studies on structural connectivity, volumetry, and anterograde tracing show that there are neuroplastic changes in (i) the colliculi (Amaral et al., 2016); (ii) the thalamus (Amaral et al., 2016; Kok & Lomber, 2017; Butler et al., 2018); and (iii) cortical regions (Beer et al., 2011; Amaral et al., 2016). For instance, Amaral and colleagues (2016) proposed that volumetric changes in subcortical structures in deaf individuals that mimic the kinds of volumetric and functional asymmetries found in the auditory cortex of those same individuals could be related to information rerouting from visual to auditory streams. These authors showed that congenitally deaf humans present hemispheric asymmetries, similar to those observed in the auditory cortex, in a number of subcortical areas. Specifically, they found the right thalamus, the right inferior colliculus, and the right lateral geniculate nucleus to be significantly larger than the left counterparts in deaf but not hearing individuals – mimicking the functional asymmetry in neuroplasticity between right and left auditory cortex in the congenitally deaf. Thus, the authors suggested that these three subcortical areas could function as a relay of visual information to the neuroplastically changed auditory cortex.

Tracing results on deaf cats also suggest that subcortical structures, and specifically the superior colliculi, may be partially involved in the passage of visual information to the auditory cortex under deafness (Butler et al., 2018). Retrograde tracers injected into the superior colliculus led to labelling in the auditory cortex (Butler et al., 2018). Although the results from Butler and colleagues (2018) suggest projections from the auditory cortex to subcortical structures, their results clearly show that the superior colliculi are connected to the auditory cortex in early deaf animals. Moreover, Kok and Lomber (2017) showed that the pattern of connections from the thalamus to the dorsal zone of the auditory cortex of early and late deaf cats remains similar to the pattern of connections of hearing cats. That is, despite the lack of auditory relay within the thalamus in congenitally deaf cats, and the potential degeneration thereof of the connections to and from the auditory cortex, no decrement in the number of connections is present following hearing loss. Possibly, the unaltered connection patterns could be due to a novel flow of visual information between thalamic and auditory areas. Lastly, it has been shown that, in hearing individuals, modulatory effects of sounds on visual performance depend on direct structural connections between auditory regions, such as the Heschl’s Gyrus and the Planum Temporale, and visual cortical areas (Beer et al., 2011). These connections may be used in congenital deafness to re-route visual information to the auditory cortex.

These data suggest a series of possible cortical and subcortical routes for visual information to reach the auditory cortex of deaf humans – some dependent on neuroplastic changes due to congenital deafness (potentially within subcortical visual and auditory relay regions) and others that perhaps are present in a normally developing brain, and that can be co-opted under deafness (potentially those involving cortico-cortical connections). Here we will test how visual information reaches the auditory cortex of the congenitally deaf, and focus on the role of these cortical and subcortical connections in the passage of visual information to the auditory cortex. We will test functional connectivity fingerprints between both cortical and subcortical regions described above (i.e., early visual cortex, Lateral Geniculate Nuclei, LGN; Medial Geniculate Nucleus, MGN; Superior Colliculi, SC; and Inferior Colliculi, IC), and the auditory cortex in the congenitally deaf, and compare those to hearing participants. First, we will look at how these regions are connected with the auditory cortex using a functional connectivity measure that takes into account the representations at the different target regions (Representational Connectivity Analysis; RCA, Nili et al., 2014). Secondly, we will compute effective connectivity between these regions using dynamic causal modelling (DCM, Friston et al., 2003). Our data shows that the superior colliculi of deaf, but not hearing, individuals feed visual information both to the early visual cortex and the inferior colliculi, and that in turn, these two feed the auditory cortex.

## Material and Methods

### Participants

Thirty-one young adults with normal or corrected to normal vision and without neurological disorders participated in this study. Sixteen were hearing individuals (13 women, group mean age = 20.1 years) and fifteen were congenitally deaf individuals (12 women, group mean age = 20.4 years). Congenitally deaf participants did not use hearing aids, had binaural hearing loss reported above 90 dB (tested frequencies were between 125 and 8,000 Hz), and were proficient in sign language. All participants read and signed an informed consent term, which was part of an experimental protocol approved by the institutional review board of Beijing Normal University Imaging Center for Brain Research. Complete datasets from five deaf and six hearing participants were excluded from further analysis due to high level of noise (i.e., head motion above 3 mm within sessions, or low fMRI signal-to-noise ratio). The remaining dataset consisted of fMRI data from ten hearing subjects (8 women, group mean age 23.1 years), and ten congenitally deaf subjects (9 women, group mean age = 23.4 years). These functional data have been used in previous publications (functional data Almeida et al, 2015; structural data Amaral et al, 2016).

### Stimuli and procedure

Participants went through four runs in the MRI scanner. Visual stimuli were presented in a blocked design with five blocks per run. Each run started with a resting period of 12 seconds and was followed by a 48-second block of stimuli. In two runs (the “Ring runs”), a random sequence of four images showing counterphase flickering (5 Hz) rings subtending 9.23°, 6.61°, 3,97°, or 1.30° of the visual angle were presented for 12 seconds each, and followed by a 12-second inter-bock-interval (IBI). Blocks in the other two runs (the “Wedge runs”) presented a random sequence of four images of counterphase flickering (5 Hz) checkerboard wedges subtending 10.50° of the visual angle that were located on the left and right parts of the screen along the azimuth plane, and upper and lower parts of the screen along the meridian plane. Each wedge stimulus was presented for 12 seconds, and a block of four consecutive wedge or ring stimuli was followed by a 12-second IBI. Participants were asked to fixate at the center of the screen for the duration of the experiment (**Figure 1A**).

**Figure1:**
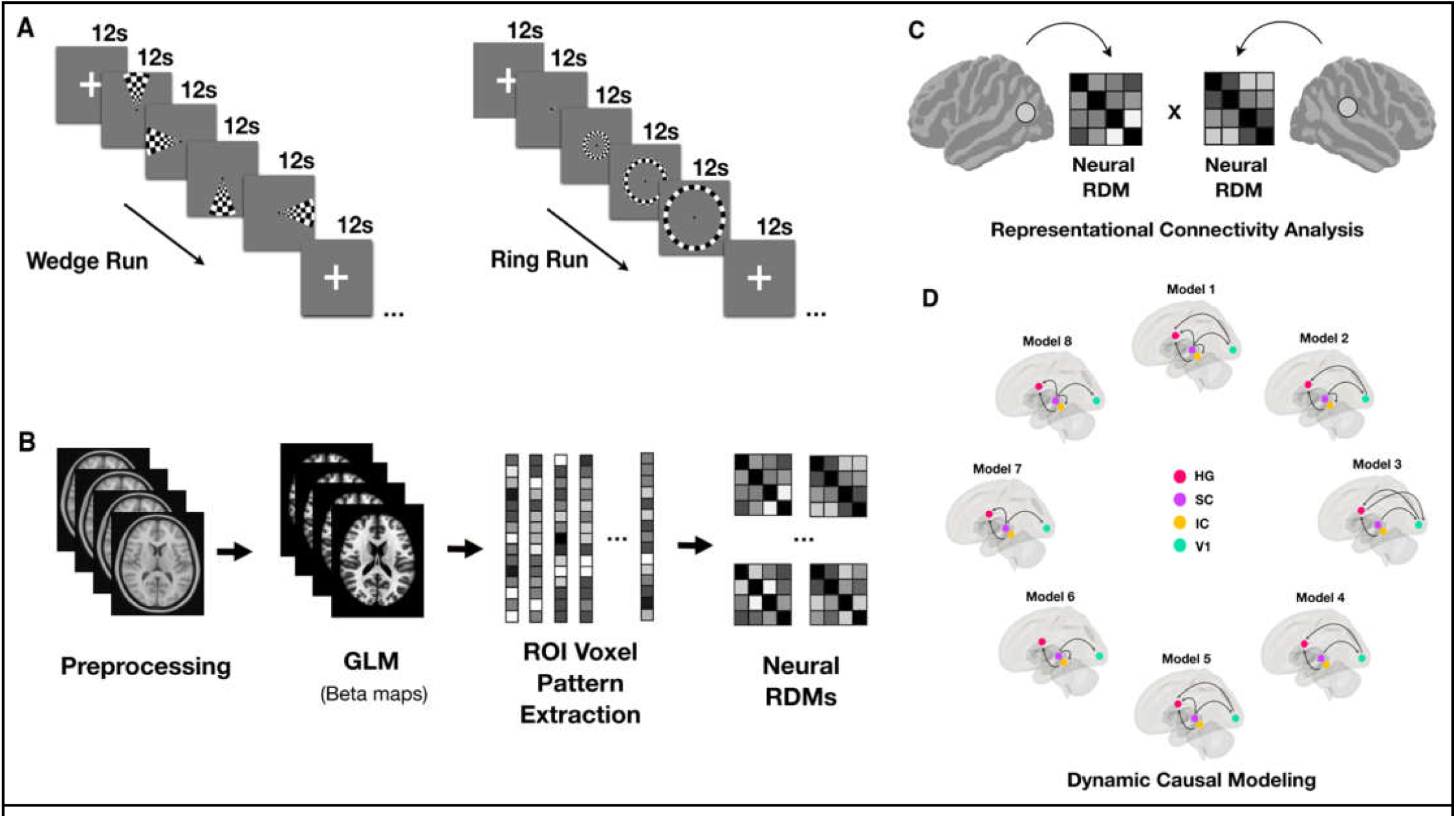
Methods outline: **A**. Experimental setup: Representations of a block of stimuli in the wedge run (left), and in the ring run (right). Stimuli were randomly presented within blocks **B**. Preprocessing pipeline followed by beta maps estimation, ROI’s feature extraction, and neural RDMs computation. **C**. Representational Connectivity Analysis scheme. **D**. Dynamic Causal Models scheme.

### MRI data acquisition

Functional and structural magnetic resonance data were acquired on a 3T Siemens Tim Trio scanner. Structural images were collected with a 3D-MPRAGE sequence (in the sagittal plane –144 slices, TR = 2530 ms, TE = 3.39 ms, FOV = 256 × 256, flip angle = 7°, voxel size = 1.33 × 1 × 1.33 mm). Functional images were collected with an EPI sequence (in axial plane – 33 slices, TR = 2000 ms, TE = 30 ms, FOV = 64 × 64, flip angle: 90°, voxel size = 3.125 × 3.125 × 4 mm, inter-slice distance = 4.6 mm).

### fMRI data pre-processing

Functional and structural data have been preprocessed on SPM12 toolbox (https://www.fil.ion.ucl.ac.uk/spm/). Preprocessing pipeline included slice-timing correction, motion regression, structural normalization into MNI 152 template, functional co-registration, linear trend removal, and high-pass filtering of 0.008 Hz.

### Selection of Regions-of-Interest (ROIs)

ROIs included right and left portions of the primary visual cortex (V1), Heschl’s gyrus (HG), Lateral Geniculate Nuclei (LGN), Medial Geniculate Nucleus (MGN), Superior Colliculi (SC), and Inferior Colliculi (IC). Right and left V1 (BA 17) regions were extracted from SPM Anatomy toolbox (Eickhoff, et al., 2005). Right and left thalamic nuclei (LGN and MGN) were extracted from the Jüelich Histological Atlas (Bürgel, et al., 1999; 2006). Right and left HG were manually segmented, in a previous study, for each participant in high resolution native space (Amaral et al., 2016), co-registered to individual functional mean image space (in native space), and normalized. Right and left superior and inferior colliculi were manually segmented in each k-slice from high resolution structural images using editing tools from FSLeyes (https://fsl.fmrib.ox.ac.uk/fsl/fslwiki). In order to define the borders of both superior and inferior colliculi we followed the description given in García-Gomar and Colleagues (2019) for anatomical identification and manual segmentation.

### Analysis

Representational connectivity, ROI-to-ROI and searchlight analysis, were performed using in-house scripts and functions from CoSMoMVPA toolbox (http://www.cosmomvpa.org/index.html) for MATLAB. Dynamic Causal Modeling analysis was performed with SPM12 toolbox.

Representational features were extracted from beta maps estimated on the first level of GLM analysis. GLM analysis estimated beta values for each voxel included in the grey matter mask created during individual structural image segmentation. A beta map was estimated for each stimulus (annuli: first, second, third and fourth; and wedge: up, down, left and right) in each run. The GLM outputs were two beta maps for each stimulus, one map per run. For each stimulus, beta maps were averaged across runs. ROI’s features were extracted from averaged beta maps, and used to calculate neural RDMs and estimate searchlight spaces.

### Representational Connectivity Analysis (RCA)

ROI-to-ROI multivariate connectivity analysis was conducted. As proposed by Kriegeskorte and colleagues (2018), Representational Similarity Analysis brings forward a powerful tool for investigating functional connections between different brain regions with a multivariate approach. In short, RCA consists of comparisons between representational (dis)similarity matrices from two different regions of interest, permitting inferences on how similar (or correlated) the information patterns encoded by both regions are, and therefore, how strongly two regions are functionally and representationally connected to each other.

In the ROI-to-ROI approach, spearman’s correlations were calculated between the seeds (right and left HG) and RDMs of regions that are part of the visual and auditory stream – focusing on subcortical regions and V1. Thus, we computed ring-based and wedge-based RDMs from the left and right LGN, right and left MGN, right and left SC, right and left IC, as well as right and left V1. These RDMs were created as means to assess how well each cortical and subcortical stage of visual and auditory pathway would correlate to representations in the Heschl’s Gyrus for deaf and hearing groups. Spearman’s correlations were calculated between left and right HG RDMs and right and left V1, LGN, MGN, SC, and IC for each subject. We used bootstrap analysis (with 10,000 resamples per test) to test the differences in correlation between HG and the remaining ROIs within and between groups.

### Dynamic Causal Modeling (DCM)

Dynamic causal modeling, first proposed by Friston et al. (2003), allows one to infer underlying neuronal state precursors of measured electrophysiological or hemodynamic data. A set of differential equations are used to describe and model the dynamic of neuronal populations hypothetically involved in the physiological phenomena, which give rise to the signals measured with fMRI scanner, for instance (Friston et al., 2003, Stephan et al., 2010). In the last stage, DCM makes use of a Bayesian framework which permits the assessment of a set of candidate models, and ranks these models according to their exceedance probability. In the present study we performed DCM analysis based on our RCA ROI-to-ROI results. We aimed to assess the dynamic interactions between visual and auditory regions in deaf subjects. The main hypothesis is that visual information might be rerouted to the auditory stream at early pre-cortical stages of visual processing/subcortical processing, namely inferior and superior colliculus. In order to test this hypothesis eight models were proposed and scored according to their exceedance probability calculated by Bayesian model selection. Models included the ROIs which showed significant correlations to HG in the RCA analysis, and had the average correlation to HG significantly different between deaf and hearing subjects. All models were based on the same four ROIs: right SC, right IC, right V1, and right HG, and all models had the SC as a doorway of visual information into the network. Models only varied in terms of existing connections and connection directionality. **Model 1:** proposes SC as a hub, and HG receives information from V1, SC, and IC. **Model 2:** proposes that SC sends information to IC, and V1. HG receives information from V1 and IC. **Model 3:** proposes that SC sends information only to V1. IC sends information only to HG. Information flows between V1 and HG bidirectionally. **Model 4:** is similar to model3, however only V1 sends information to HG. **Model 5:** is similar to model 1, however SC is not connected to IC. **Model 6:** suggests that SC sends information to V1, and SC. IC sends information to HG. **Model 7:** is similar to model 6, however SC is not connected to IC, but SC is directly connected to HG. Finally, **Model 8:** is similar to model 1, however V1 and HG are not connected between them. Differences between models rely only on network extrinsic connectivity (**Figure 4A**).

## Results

### Representational Connectivity Analysis

In order to test whether cortical and subcortical visual and auditory regions represent visual information similarly to the HG, we performed bootstrap analysis (with 10,000 resamples) on the correlation values between HG (left and right) and each of the other ROI RDMs. Main results are reported here, and comprise group mean Spearman’s coefficient (M), standard error of the mean (SEM), confidence interval (CI), and significant FDR-corrected p-values. Tables with expanded analysis results within groups are reported in Supplementary Material - Section 1.

#### Ring stimuli

RCA analysis compared HG RDMs to RDMs from right and left V1, LGN, MGN, SC, and IC. Focusing on ipsilateral connections, correlations were tested within ROI RDMs from the same side of the target RDM (i.e. right-right or left-left). Results for the deaf group showed a significant correlation between right HG RDM and right SC RDM (M_rho_=0.57 ± SEM=0.11, CI=[0.39, 0.71], p<0.001), right IC RDM (M_rho_=0.47 ± SEM=0.08, CI=[0.35, 0.59], p<0.001), and right V1 RDM (M_rho_=0.47 ± SEM=0.12, CI=[0.29, 0.64], p<0.001). There was no significant correlation between right HG RDM and right MGN (M_rho_=0.23 ± SEM=0.17, CI=[-0.05, 0.46], p=0.101) or right LGN RDMs (M_rho_=0.06 ± SEM=0.15, CI=[-0.15, 0.28], p=0.333) (**Figure 2A**). No significant correlations were found between hearing right HG RDMs and other ROI RDMs. Between group comparisons indicated that right SC (M_Deaf_ - M_Hearing_=0.49, CI=[0.15, 0.81], p=0.028,), right IC (M_Deaf_ - M_Hearing_=0.38, CI=[0.05, 0.72], p=0.045), and right V1 (M_Deaf_ - M_Hearing_=0.47, CI=[0.11, 0.82], p=0.033) are significantly differently correlated to HG RDMs between deaf and hearing groups. On the left side, only left V1 RDM showed a significantly positive correlation to left HG RDM for the deaf group (M_rho_=0.36 ± SEM=0.19, CI=[-0.15, 0.28], p=0.043). No significant differences between groups were found on the left side (**Figure 2B**).

**Figure 2:**
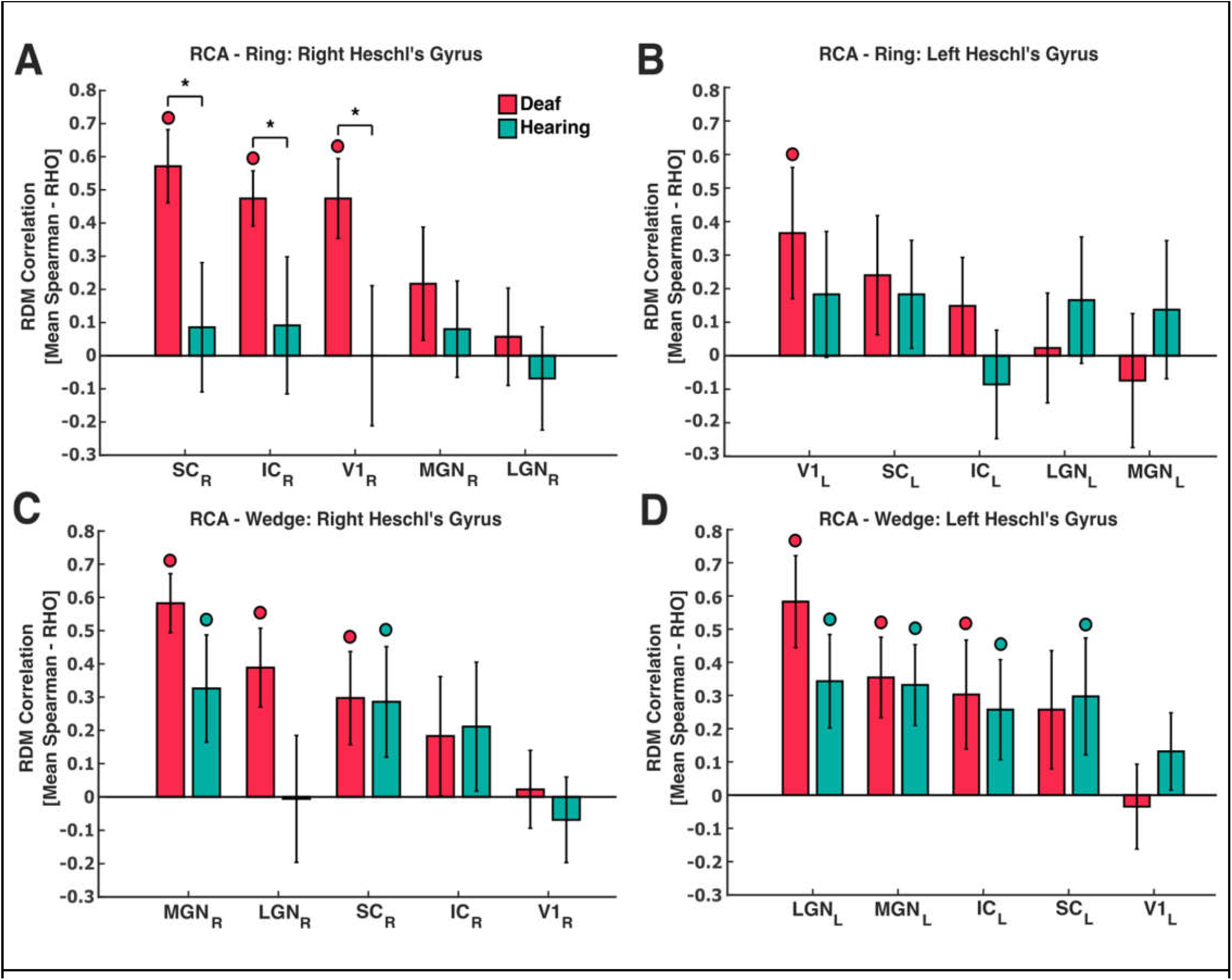
Representational connectivity analysis. **A + B**. Spearman’s group average correlations and SEM between right (A) and left (B) HG and five ROIs from visual and auditory streams for deaf (red bars) and hearings (green bars) for the Ring condition (SC = Superior Colliculus, IC = Inferior Colliculus, LGN = Lateral Geniculate Nucleus, MGN = Medial Geniculate Nucleus, V1 = Early Visual Area). **C + D**. Spearman’s group average correlations and SEM between right (C) and left (D) HG and five ROIs from visual and auditory streams for deaf (red bars) and hearings (green bars) for the Wedge condition. Red or green colored “**o**” indicates a significant correlation within each group (FDR corrected for multiple comparisons), and “**\***” indicates significant differences in correlation between groups (FDR corrected for multiple comparisons).

#### Wedge stimuli

RCA ROI-to-ROI analysis compared HG RDMs to RDMs from V1, LGN, MGN, SC, and IC. Results showed significant correlations between right deaf HG RDM and right MGN (M_rho_=0.58 ± SEM=0.09, CI=[0.44, 0.70], p<0.001), right LGN (M_rho_=0.39 ± SEM=0.12, CI=[0.21, 0.56], p<0.001), and right SC (M_rho_=0.30 ± SEM=0.14, CI=[0.08, 0.50], p=0.011). Hearing right HG RDM significantly correlated to right MGN (M_rho_=0.32 ± SEM=0.16, CI=[0.08, 0.56], p=0.039), and right SC (M_rho_=0.29 ± SEM=0.16, CI=[0.02, 0.52], p=0.039). No between group differences were found significant between right HG and other ROIs connectivity (**Figure 3C**). On the left side, deaf left HG RDM significantly correlated to left LGN (M_rho_=0.58 ± SEM=0.14, CI=[0.36, 0.77], p<0.001), left MGN (M_rho_=0.35 ± SEM=0.12, CI=[0.17, 0.53], p<0.001), and left IC (M_rho_=0.30 ± SEM=0.16, CI=[0.05, 0.54], p=0.028). Hearing left HG RDM significantly correlated to LGN (M_rho_=0.34 ± SEM=0.14, CI=[0.13, 0.55], p=0.004), MGN (M_rho_=0.33 ± SEM=0.12, CI=[0.15, 0.50], p<0.001), IC (M_rho_=0.26 ± SEM=0.15, CI=[0.03, 0.47], p=0.034) and SC (M_rho_=0.30 ± SEM=0.17, CI=[0.03, 0.55], p=0.034). No between group differences were found significant in between left HG and other ROIs connectivity (**Figure 2D**).

**Figure 3:**
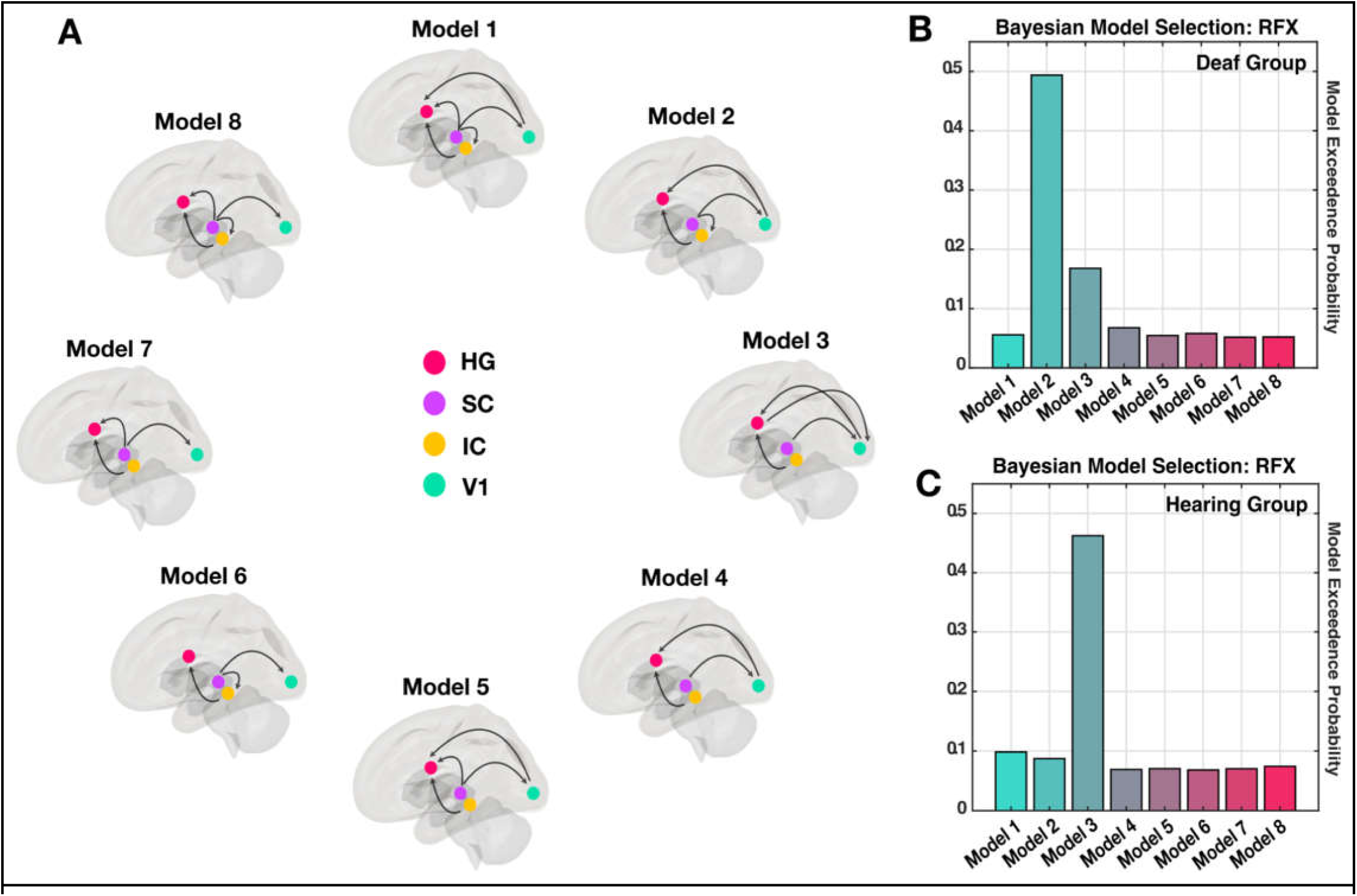
Dynamic causal modeling. **A**. Eight DCM candidate models. All models included four nodes: right Heschl’s gyrus (HG, pink), right Superior Colliculus (SC, purple), right Inferior Colliculus (IC, yellow), and right early visual area (V1, green). Black arrows indicate the intrinsic dynamic of each model and the direction the visual information flows between nodes **B**. Results from Bayesian Model Selection with Random Effects for the deaf group. Exceedance probability of each model was plotted, and suggests model 2 is the strongest candidate model for the deaf group. **C**. Results from Bayesian Model Selection with Random Effects for the hearing group. Exceedance probability of each model was plotted, and suggests model 3 is the strongest candidate model for the hearing group.

### Dynamic Causal Modeling (DCM)

In the RCA results, the right SC, right IC, and right V1 were found to significantly correlate to right HG of deaf individuals, but not hearing individuals, when presented with annuli stimuli. Because of that we further created eight different candidate models in order to assess the directionality of connections between nodes using DCM. All models were based on the same four ROIs: right SC, right IC, right V1, and right HG (**Figure 3A**).

A first level General Linear Model analysis was defined using all four annuli stimuli. Annuli stimuli were set as the modulatory inputs of the eight network models. Time series from right SC, right IC, right V1, and right HG were then extracted. DCM models were set including bilinear modulatory effects, and no stochastic effect. After specifying all models for hearing and deaf subjects, the eight models were compared using a Bayesian model selection approach. A random effects approach was used as a second level inference for both groups. Model exceedance probabilities for deaf group were: Model 1 = 0.055; Model 2 = 0.493; Model 3 = 0.167; Model 4 = 0.067; Model 5 = 0.054; Model 6 = 0.057; Model 7 = 0.051, and Model 8 = 0.052 (**Figure 3B**). Model exceedance probabilities for hearing group were: Model 1 = 0.098; Model 2 = 0.087; Model 3 = 0.462; Model 4 = 0.069; Model 5 = 0.070; Model 6 = 0.068; Model 7 = 0.070, and Model 8 = 0.074 (**Figure 3B**). That is, for hearing individuals we show the typical separation between auditory and visual streams, with (bidirectional) connections between these two sensory streams happening exclusively at the cortical level, whereas for congenitally deaf individuals, auditory cortex receives visual information not only from visual cortical regions, but importantly from the superior colliculi via the inferior colliculi.

## Discussion

The present study aimed to investigate which cortical and subcortical areas could be involved in rerouting visual information to the early auditory cortex of congenitally deaf individuals. For that, we evaluated the plausibility of a model of visual information processing in deaf individuals, taking into account the possible connections between visual and auditory systems at both the cortical and subcortical levels. First, we searched for potential node candidates using RCA analysis over both eccentricity and polar angle representations. Our main results from this analysis showed that the right auditory cortex (the right HG) represents visual eccentricity information similarly to the right primary visual cortex, right superior colliculus and right inferior colliculus. Notably, this is true only in the congenitally deaf group. Secondly, we aimed at characterizing the dynamics of connections between these nodes in both deaf and hearing groups. Using dynamic causal modelling, we show that early visual cortex streams visual information to the auditory cortex in both hearing and deaf individuals as previously shown (Beer et al., 2011). More importantly, however, we found an up-to-now novel route that may carry visual information to the primary auditory cortex and that is exclusive to congenitally deaf individuals. Specifically, we found a subcortical connection that goes from the superior colliculi (via the inferior colliculi) to the auditory cortex of deaf individuals.

Multivariate connectivity analysis showed that the HG represents visual information of eccentricity similarly to subcortical auditory and visual regions in deaf but not hearing individuals. We used right and left HG as seeds, and targeted other cortical and subcortical visual and auditory streams ROIs, namely the superior and inferior colliculus, the lateral and medial geniculate nucleus and early visual cortex. For experimental conditions where we showed participants annuli stimuli, RCA results showed that right superior colliculus, right inferior colliculus and right V1 were more strongly connected to the right HG of deaf, but not hearing individuals. No other between group comparisons showed significant differences of representational connectivity (in the ROI-to-ROI or the whole-brain searchlight). That is, visual representations in right superior colliculus, right inferior colliculus and right V1 were similar to the visual representations in the HG of the deaf but not of the hearings, which is consistent with some of the effects presented in the literature (e.g., Almeida et al., 2015).

Although some of the between group comparisons for the experimental condition where we presented wedge stimuli to our participants were close to significance – for instance, stronger RCA values between the right HG and the right LGN for the deaf group when compared to the hearing group were significant at an uncorrected p-value (uncorrected-p = 0.035, FDR-corrected-p = 0.134) – we did not include data from this condition in further analysis. A possible reason why we failed to find significant correlations between the representations in HG and other brain areas (e.g., V1) is that polar angle representations in HG were shown to be limited to the horizontal plane (Almeida et al., 2015; 2018). Therefore, while visual areas (e.g., V1) fully represent polar angle visual information in vertical and horizontal planes, the HG of deaf individuals only partially represents this visual information (i.e., only on the horizontal plane), possibly leading to weaker correlations between the representational content within those regions.

In order to investigate the dynamic functional interaction among the regions described above, we tested different possible pathways of visual information from visual to auditory regions with dynamic causal modelling. In a previous study, Klinge and colleagues (2010) used DCM to investigated how auditory information reached the visual cortex under blindness. They focused on thalamocortical and corticocortical connections, and used dynamic connectivity (dynamic causal modelling; DCM) between early visual cortex, early auditory cortex, and thalamic medial geniculate nucleus (MGN), and showed that MGN works as a redistributor of auditory information between the early visual and auditory cortices in blind individuals. These results highlighted the importance of subcortical structures in re-routing sensory information to the sensory-deprived cortex. Similarly, here, we employed DCM to investigate the role of subcortical and cortical structures from visual and auditory streams in re-routing the flow of visual information to the auditory cortex of deaf individuals. We proposed eight different dynamic models to be tested with DCM. Each model proposed different dynamics of connections between the four regions found to be connected to the right HG in deaf individuals.

As hypothesized, Model 3, which describes what is considered the dynamic of visual information in hearing individuals (Beer et al., 2011), was the model which stands up with the highest exceedance probability for the hearing group. Notably, Model 2 was the model with the highest exceedance probability for the deaf group. Model 2 suggests a novel route of visual information from right superior colliculus to right inferior colliculus heading to deaf early auditory cortex. Moreover, and similarly to Model 3, Model 2 also suggests a flow of visual information from right early visual area to right auditory area. In summary, our DCM analysis from our eight candidate models resulted in a winning model for the hearing group, and a different model for the deaf group. The winning model for deaf individuals suggests that visual information reaches the early auditory area of deaf individuals from two different sources: corticocortical connections between the right early visual and the right early auditory regions, and from a rerouting of visual information arising from right intercollicular connections from SC to IC.

These results suggest then that visual information reaches early auditory cortex via i) re-routing of visual information from SC to IC, and from there to HG, and ii) corticocortical projections from V1 to HG. Projections from the SC to the ipsilateral IC have been exhaustively described in non-human animal studies using retrograde and anterograde horseradish peroxidase tracer (Adams, 1980; Covey et. al.,1987; and Coleman and Clerici, 1987). Moreover, it has been reported that a number of cells in the IC are capable of responding exclusively to visual stimuli (Porter et al., 2007; Bulkin and Groh, 2012; Gruters and Groh, 2012). Thus, extant literature supports the connection proposed herein, and the potential role of the SC and IC in re-routing visual information to the HG. Regarding the corticocortical connections between early visual cortex and the HG, Beer and colleagues (2011) reported a direct structural connection between early auditory cortex and early visual cortex in healthy humans. Although these data on the connectivity between IC and SC and between early visual and auditory cortices were found in hearing individuals, it is nevertheless highly probable that neuroplasticity co-opted this, and strengthened an already existing set of connections between visual and auditory streams. In fact, Amaral and colleagues (2016) showed that the right IC of deaf individuals was larger than its contralateral – suggesting perhaps an increment of connections in IC due to its recruitment in visual information processing.

A possible limitation of our model for deaf individuals could be related to the selection of the doorway for visual information into the system. All eight proposed models assume visual information enters the network via SC. Nevertheless, LGN is the major route of visual information to the cortex. We selected the SC because the SC was highly correlated with HG in our RCA analysis, and thus seemed to be the best distributor of visual information to the network under deafness.

## Conclusion

The present study investigated how visual information is re-routed to the auditory cortex of congenitally deaf individuals. We suggest a mechanistic model of how visual information reaches the early auditory cortex of congenitally deaf individuals, whereby visual information reaches the early auditory cortex of deaf individuals via an intercollicular rerouting of visual information from superior to inferior colliculus that is unique to deaf individuals, and a corticocortical functional projection from the early visual area to the early auditory area that is also present in hearing individuals. That is, neuroplastic changes due to congenital deafness led to a rewiring of the system at the subcortical level in order to allow for congenitally sensorially deprived cortices to receive other sensorial information.

## Acknowledgements

This research is supported by an FCT grant PTDC/PSI-GER/30757/2017 AND Programa COMPETE Portugal. JA, GMG, and ZT are supported by funds from the European Research Council (ERC) under the European Union’s Horizon 2020 research and innovation programme Starting Grant number 802553; “ContentMAP” attributted to JA. FF is supported by the National Natural Science Foundation of China (31930053).

## Supplementary Material

### RCA: ROI-to-ROI results

Representational Connectivity Analysis: correlations between right and left HG and ipsi-lateral V1, superior colliculus (SC), inferior colliculus (IC), lateral geniculate nucleus (LGN), and medial geniculate nucleus (MGN) for eccentricity and polar angle conditions.

**Table 1.**
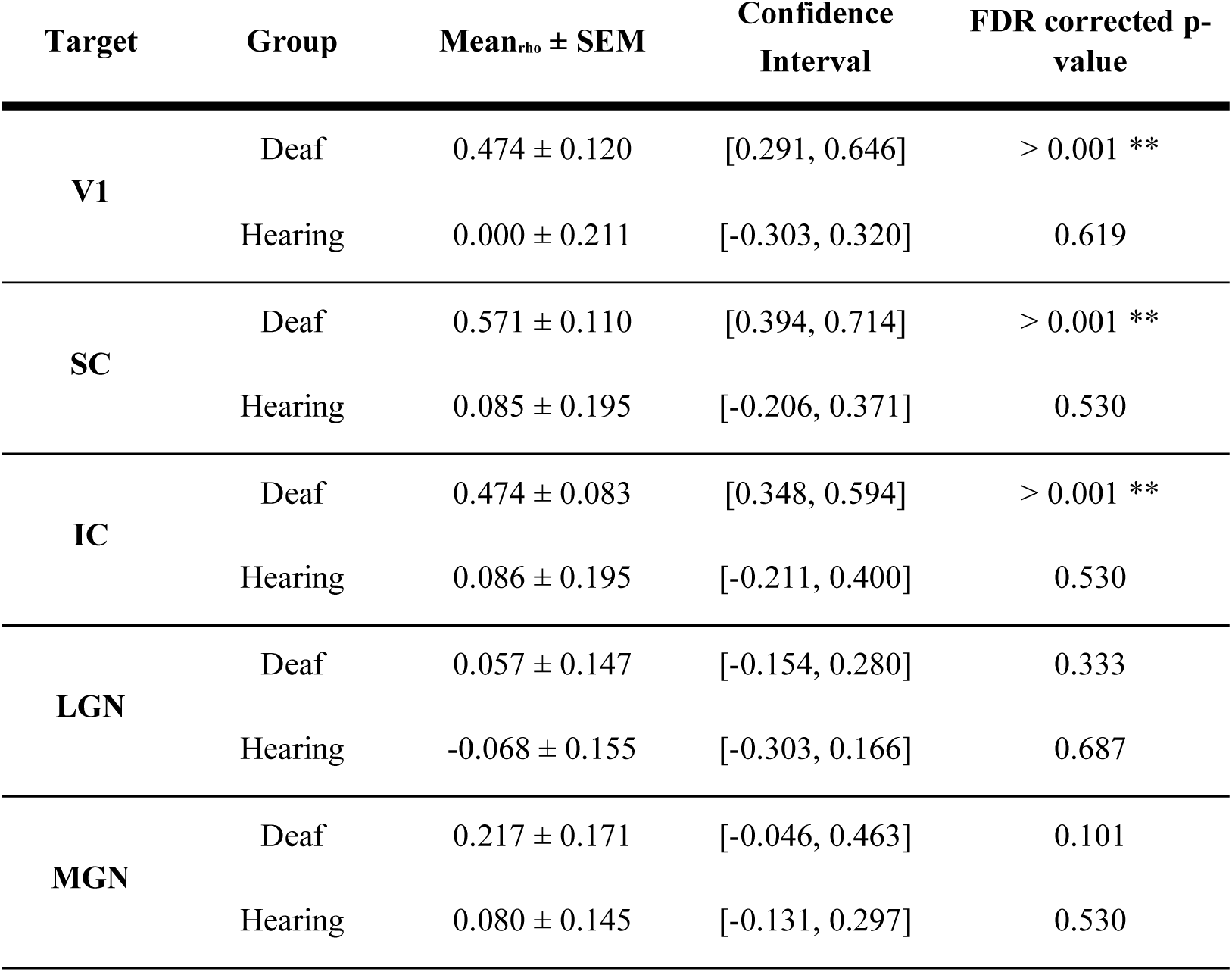
Correlations between right HG and regions of interest for hearing and deaf groups in eccentricity condition. ** = significant.

**Table 2.**
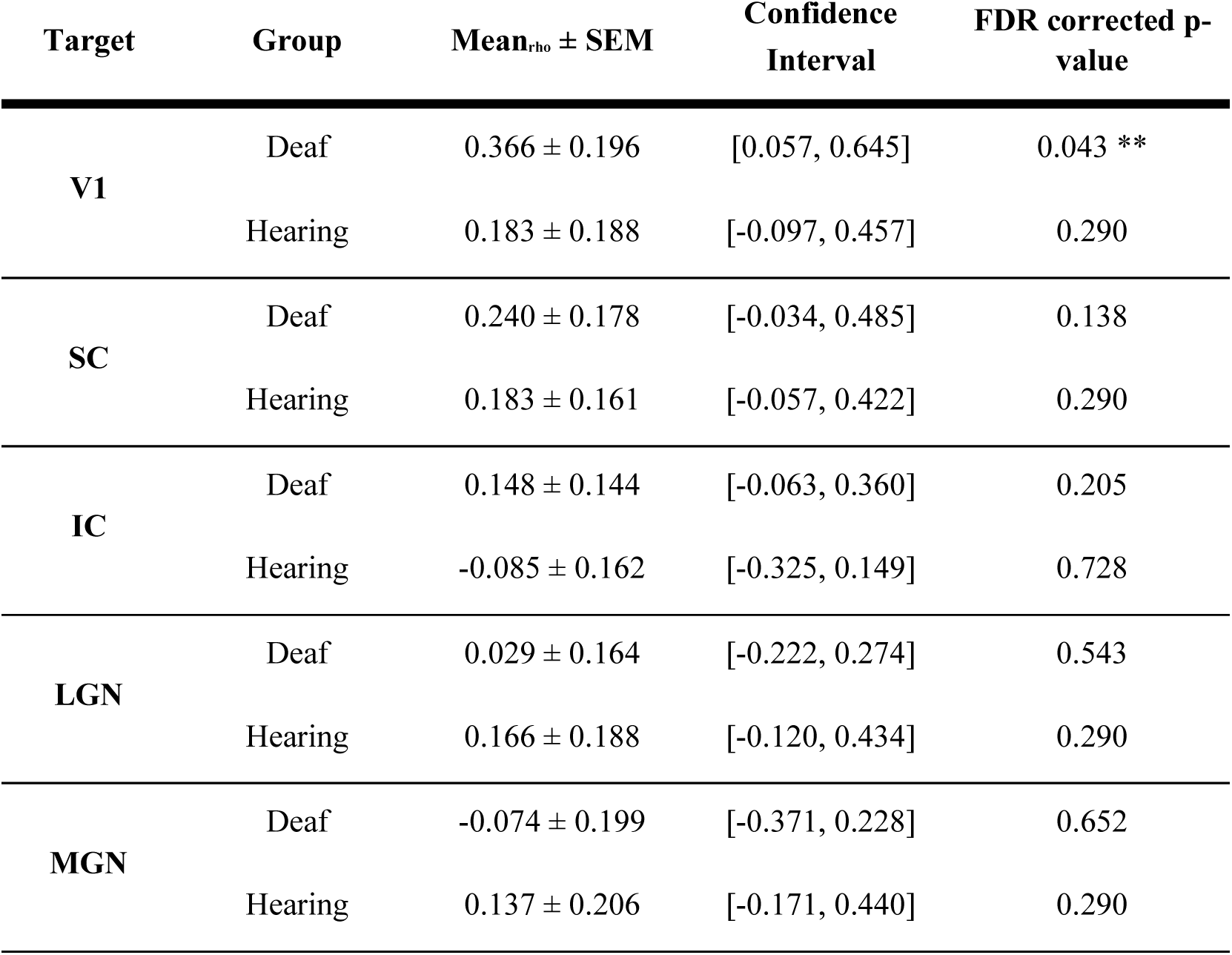
Correlations between left HG and regions of interest for hearing and deaf groups in eccentricity condition. ** = significant.

**Table 3.**
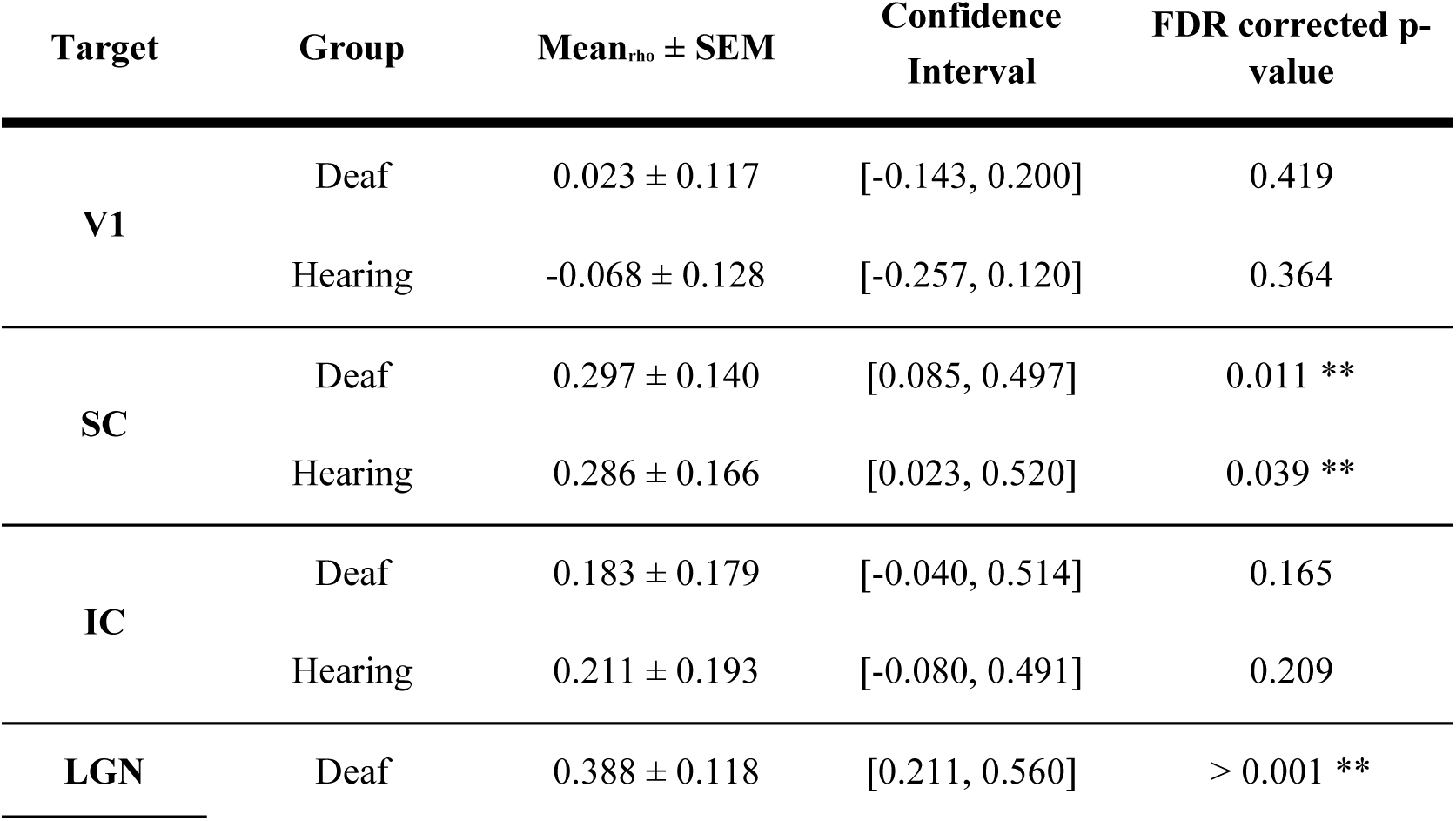
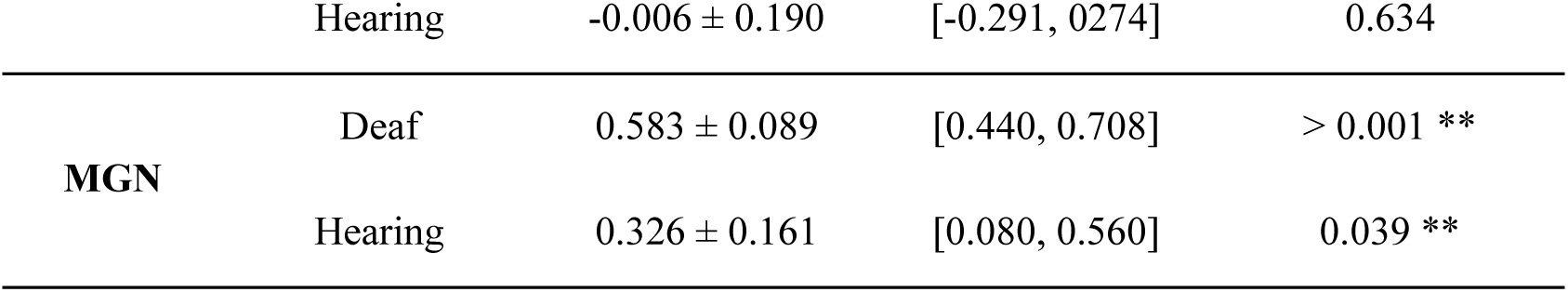
Correlations between right HG and regions of interest for hearing and deaf groups in polar angle condition. ** = significant.

| Target | Group | Mean<sub>rho</sub> ± SEM | Confidence Interval | FDR corrected p-value |
| --- | --- | --- | --- | --- |
| V1 | Deaf | 0.023 ± 0.117 | [-0.143, 0.200] | 0.419 |
|  | Hearing | -0.068 ± 0.128 | [-0.257, 0.120] | 0.364 |
| SC | Deaf | 0.297 ± 0.140 | [0.085, 0.497] | 0.011 ** |
|  | Hearing | 0.286 ± 0.166 | [0.023, 0.520] | 0.039 ** |
| IC | Deaf | 0.183 ± 0.179 | [-0.040, 0.514] | 0.165 |
|  | Hearing | 0.211 ± 0.193 | [-0.080, 0.491] | 0.209 |
| LGN | Deaf | 0.388 ± 0.118 | [0.211, 0.560] | > 0.001 ** |
|  | Hearing | -0.006 ± 0.190 | [-0.291, 0.274] | 0.634 |
| **MGN** | Deaf | 0.583 ± 0.089 | [0.440, 0.708] | > 0.001 ** |
|  | Hearing | 0.326 ± 0.161 | [0.080, 0.560] | 0.039 ** |

**Table 4.** Correlations between left HG and regions of interest for hearing and deaf groups in polar angle condition. *= marginally significant, and ** = significant.

| Target | Group | Mean<sub>rho</sub> ± SEM | Confidence Interval | FDR corrected p-value |
| --- | --- | --- | --- | --- |
| **V1** | Deaf | -0.034 ± 0.127 | [0.222, 0.148] | 0.623 |
|  | Hearing | 0.131 ± 0.116 | [-0.028, 0.314] | 0.105 |
| **SC** | Deaf | 0.257 ± 0.178 | [-0.011, 0.534] | 0.056 * |
|  | Hearing | 0.297 ± 0.176 | [-0.063, 0.514] | 0.034 ** |
| **IC** | Deaf | 0.303 ± 0.164 | [0.051, 0.543] | 0.028 ** |
|  | Hearing | 0.257 ± 0.251 | [0.034, 0.474] | 0.034 ** |
| **LGN** | Deaf | 0.583 ± 0.138 | [0.366, 0.771] | > 0.001 ** |
|  | Hearing | 0.343 ± 0.141 | [0.131, 0.554] | 0.004 ** |
| **MGN** | Deaf | 0.354 ± 0.121 | [0.177, 0.531] | > 0.001 ** |
|  | Hearing | 0.331 ± 0.122 | [0.148, 0.503] | > 0.001 ** |

## References

Alencar, C. D., Butler, B. E., & Lomber, S. G. (2019). What and how the deaf brain sees. Journal of cognitive neuroscience, 31(8), 1091–1109.

Adams, J. C. (1980). Crossed and descending projections to the inferior colliculus. Neuroscience Letters, 19(1), 1–5. doi:10.1016/0304-3940(80)90246-3

Amaral, L. & Almeida, J. (2015) Neuroplasticity in congenital deaf humans. Revista Portuguesa de Psicologia, 44, 39–45

Amaral, L., Ganho-Ávila, A., Osório, A., Soares, M. J., He, D., Chen, Q., Almeida, J. (2016). Hemispheric asymmetries in subcortical visual and auditory relay structures in congenital deafness. European Journal of Neuroscience, 44(6), 2334–2339. doi:10.1111/ejn.13340

Almeida, J., He, D., Chen, Q., Mahon, B. Z., Zhang, F., Gonçalves, Ó. F., Bi, Y. (2015). Decoding Visual Location From Neural Patterns in the Auditory Cortex of the Congenitally Deaf. Psychological Science, 26(11), 1771–1782. doi:10.1177/0956797615598970

Almeida, J., Nunes, G., Marques, J. F., & Amaral, L. (2018). Compensatory plasticity in the congenitally deaf for visual tasks is restricted to the horizontal plane. Journal of Experimental Psychology: General, 147(6), 924.

Beer, A. L., Plank, T., & Greenlee, M. W. (2011). Diffusion tensor imaging shows white matter tracts between human auditory and visual cortex. Experimental Brain Research, 213(2-3), 299–308.doi:10.1007/s00221-011-2715-y

Bell, L., Wagels, L., Neuschaefer-Rube, C., Fels, J., Gur, R. E., & Konrad, K. (2019). The Cross-Modal Effects of Sensory Deprivation on Spatial and Temporal Processes in Vision and Audition: A Systematic Review on Behavioral and Neuroimaging Research since 2000. Neural plasticity.

Benetti, S., van Ackeren, M. J., Rabini, G., Zonca, J., Foa, V., Baruffaldi, F., & Collignon, O. (2017). Functional selectivity for face processing in the temporal voice area of early deaf individuals. Proceedings of the National Academy of Sciences, 114(31), E6437–E6446.

Bola, Ł., Zimmermann, M., Mostowski, P., Jednoróg, K., Marchewka, A., Rutkowski, P., & Szwed, M. (2017). Task-specific reorganization of the auditory cortex in deaf humans. Proceedings of the National Academy of Sciences, 114(4), E600–E609.

Bosworth, R. G., Petrich, J. A., & Dobkins, K. R. (2013). Effects of attention and laterality on motion and orientation discrimination in deaf signers. Brain and cognition, 82(1), 117–126.

Bottari, D., Caclin, A., Giard, M. H., & Pavani, F. (2011). Changes in early cortical visual processing predict enhanced reactivity in deaf individuals. PloS one, 6(9), e25607.

Bottari, D., Heimler, B., Caclin, A., Dalmolin, A., Giard, M. H., & Pavani, F. (2014). Visual change detection recruits auditory cortices in early deafness. Neuroimage, 94, 172–184.

Bulkin, D. A., & Groh, J. M. (2012b). Distribution of visual and saccade related information in the monkey inferior colliculus. Frontiers in Neural Circuits, 6.doi:10.3389/fncir.2012.00061

Bürgel, U., Amunts, K., Hoemke, L., Mohlberg, H., Gilsbach, J. M., & Zilles, K. (2006). White matter fiber tracts of the human brain: Three-dimensional mapping at microscopic resolution, topography and intersubject variability. NeuroImage, 29(4), 1092–1105.doi:10.1016/j.neuroimage.2005.08.040

Bürgel, U., Schormann, T., Schleicher, A., & Zilles, K. (1999). Mapping of Histologically Identified Long Fiber Tracts in Human Cerebral Hemispheres to the MRI Volume of a Reference Brain: Position and Spatial Variability of the Optic Radiation. NeuroImage, 10(5), 489–499.doi:10.1006/nimg.1999.0497

Butler, B. E., Sunstrum, J. K., & Lomber, S. G. (2018). Modified Origins of Cortical Projections to the Superior Colliculus in the Deaf: Dispersion of Auditory Efferents. The Journal of Neuroscience, 38(16), 4048–4058. doi:10.1523/jneurosci.2858-17.2018

Cardin, V., Orfanidou, E., Rönnberg, J., Capek, C. M., Rudner, M., & Woll, B. (2013). Dissociating cognitive and sensory neural plasticity in human superior temporal cortex. Nature communications, 4(1), 1–5.

Codina, C. J., Pascalis, O., Baseler, H. A., Levine, A. T., & Buckley, D. (2017). Peripheral visual reaction time is faster in deaf adults and British sign language interpreters than in hearing adults. Frontiers in psychology, 8, 50.

Coleman, J. R., & Clerici, W. J. (1987). Sources of projections to subdivisions of the inferior colliculus in the rat. The Journal of Comparative Neurology, 262(2), 215–226. doi:10.1002/cne.902620204

Collignon, O., Voss, P., Lassonde, M., & Lepore, F. (2009). Cross-modal plasticity for the spatial processing of sounds in visually deprived subjects. Experimental brain research, 192(3), 343–358.

Covey, E., Hall, W. C., & Kobler, J. B. (1987). Subcortical connections of the superior colliculus in the mustache bat, Pteronotus parnellii. The Journal of Comparative Neurology, 263(2), 179–197.doi:10.1002/cne.902630203

Dye, M. W., Hauser, P. C., & Bavelier, D. (2009). Is visual selective attention in deaf individuals enhanced or deficient? The case of the useful field of view. PloS one, 4(5), e5640.

Eickhoff, S. B., Stephan, K. E., Mohlberg, H., Grefkes, C., Fink, G. R., Amunts, K., & Zilles, K. (2005). A new SPM toolbox for combining probabilistic cytoarchitectonic maps and functional imaging data. NeuroImage, 25(4), 1325–1335.doi:10.1016/j.neuroimage.2004.12.034

Fine, I., Finney, E. M., Boynton, G. M., & Dobkins, K. R. (2005). Comparing the effects of auditory deprivation and sign language within the auditory and visual cortex. Journal of cognitive neuroscience, 17(10), 1621–1637.

Finney, E. M., Clementz, B. A., Hickok, G., & Dobkins, K. R. (2003). Visual stimuli activate auditory cortex in deaf subjects: evidence from MEG. Neuroreport, 14(11), 1425–1427.

Finney, E. M., Fine, I., & Dobkins, K. R. (2001). Visual stimuli activate auditory cortex in the deaf. Nature neuroscience, 4(12), 1171–1173.

Friston, K. J., Harrison, L., & Penny, W. (2003). Dynamic causal modelling. Neuroimage, 19(4), 1273–1302.

García-Gomar, M. G., Strong, C., Toschi, N., Singh, K., Rosen, B. R., Wald, L. L., & Bianciardi, M. (2019). In vivo Probabilistic Structural Atlas of the Inferior and Superior Colliculi, Medial and Lateral Geniculate Nuclei and Superior Olivary Complex in Humans Based on 7 Tesla MRI. Frontiers in Neuroscience, 13.doi:10.3389/fnins.2019.00764

Gruters, K. G., & Groh, J. M. (2012). Sounds and beyond: multisensory and other non-auditory signals in the inferior colliculus. Frontiers in neural circuits, 6, 96. doi:10.3389/fncir.2012.00096

Hauthal, N., Sandmann, P., Debener, S., & Thorne, J. D. (2013). Visual movement perception in deaf and hearing individuals. Advances in cognitive psychology, 9(2), 53.

Herbin, M., Repérant, J., & Cooper, H. M. (1994). Visual system of the fossorial molelemmings, Ellobius talpinusandEllobius lutescens. Journal of Comparative Neurology, 346(2), 253–275.doi:10.1002/cne.903460206

Hribar, M., Šuput, D., Battelino, S., & Vovk, A. (2020). Review Article: Structural brain alterations in prelingually deaf. NeuroImage, 117042. doi:10.1016/j.neuroimage.2020.117042

Hunt, D. L., Yamoah, E. N., & Krubitzer, L. (2006). Multisensory plasticity in congenitally deaf mice: how are cortical areas functionally specified?. Neuroscience, 139(4), 1507–1524.

Itaya, S. K., & Van Hoesen, G. W. (1982). Retinal innervation of the inferior colliculus in rat and monkey. Brain Research, 233(1), 45–52. doi:10.1016/0006-8993(82)90928-3

Klinge, C., Eippert, F., Roder, B., & Buchel, C. (2010). Corticocortical Connections Mediate Primary Visual Cortex Responses to Auditory Stimulation in the Blind. Journal of Neuroscience, 30(38), 12798–12805. doi:10.1523/jneurosci.2384-10.2010

Kok, M. A., & Lomber, S. G. (2017). Origin of the thalamic projection to dorsal auditory cortex in hearing and deafness. Hearing Research, 343, 108–117. doi:10.1016/j.heares.2016.05.013

Kriegeskorte, N., Mur, M., & Bandettini, P. A. (2008). Representational similarity analysis-connecting the branches of systems neuroscience. Frontiers in systems neuroscience, 2, 4.

Land, R., Baumhoff, P., Tillein, J., Lomber, S. G., Hubka, P., & Kral, A. (2016). Cross-modal plasticity in higher-order auditory cortex of congenitally deaf cats does not limit auditory responsiveness to cochlear implants. Journal of Neuroscience, 36(23), 6175–6185.

Levänen, S., & Hamdorf, D. (2001). Feeling vibrations: enhanced tactile sensitivity in congenitally deaf humans. Neuroscience letters, 301(1), 75–77.

Loke, H. W., & Song, S. (1991). Central and peripheral visual processing in hearing and nonhearing individuals. Bulletin of the Psychonomic Society, 29(5), 437–440.doi:10.3758/bf03333964

Lomber, S. G., Meredith, M. A., & Kral, A. (2010). Cross-modal plasticity in specific auditory cortices underlies visual compensations in the deaf. Nature Neuroscience, 13(11), 1421–1427.doi:10.1038/nn.2653

Merabet, L. B., & Pascual-Leone, A. (2009). Neural reorganization following sensory loss: the opportunity of change. Nature Reviews Neuroscience, 11(1), 44–52. doi:10.1038/nrn2758

Meredith, M. A., & Lomber, S. G. (2011). Somatosensory and visual crossmodal plasticity in the anterior auditory field of early-deaf cats. Hearing research, 280(1-2), 38–47.

Meredith, M. A., Keniston, L. P., & Allman, B. L. (2012). Multisensory dysfunction accompanies crossmodal plasticity following adult hearing impairment. Neuroscience, 214, 136–148.

Meredith, M. A., Clemo, H. R., Corley, S. B., Chabot, N., & Lomber, S. G. (2016). Cortical and thalamic connectivity of the auditory anterior ectosylvian cortex of early-deaf cats: Implications for neural mechanisms of crossmodal plasticity. Hearing research, 333, 25–36.

Nava, E., Bottari, D., Zampini, M., & Pavani, F. (2008). Visual temporal order judgment in profoundly deaf individuals. Experimental brain research, 190(2), 179–188.

Neville, H. J., & Lawson, D. (1987b). Attention to central and peripheral visual space in a movement detection task: an event-related potential and behavioral study. II. Congenitally deaf adults. Brain Research, 405(2), 268–283. doi:10.1016/0006-8993(87)90296-4

Nili, H., Wingfield, C., Walther, A., Su, L., Marslen-Wilson, W., & Kriegeskorte, N. (2014). A Toolbox for Representational Similarity Analysis. PLoS Computational Biology, 10(4), e1003553.doi:10.1371/journal.pcbi.1003553

Porter, K. K., Metzger, R. R., & Groh, J. M. (2007). Visual-and saccade-related signals in the primate inferior colliculus. Proceedings of the National Academy of Sciences, 104(45), 17855–17860.doi:10.1073/pnas.0706249104

Sadato, N., Yamada, H., Okada, T., Yoshida, M., Hasegawa, T., Matsuki, K. I., … & Itoh, H. (2004). Age-dependent plasticity in the superior temporal sulcus in deaf humans: a functional MRI study. BMC neuroscience, 5(1), 1–6.

Scott, G. D., Karns, C. M., Dow, M. W., Stevens, C., & Neville, H. J. (2014). Enhanced peripheral visual processing in congenitally deaf humans is supported by multiple brain regions, including primary auditory cortex. Frontiers in Human Neuroscience, 8.doi:10.3389/fnhum.2014.00177

Seymour, J. L., Low, K. A., Maclin, E. L., Chiarelli, A. M., Mathewson, K. E., Fabiani, M., & Dye, M. W. (2017). Reorganization of neural systems mediating peripheral visual selective attention in the deaf: An optical imaging study. Hearing research, 343, 162–175.

Shiell, M. M., Champoux, F., & Zatorre, R. J. (2014). Enhancement of Visual Motion Detection Thresholds in Early Deaf People. PLoS ONE, 9(2), e90498. doi:10.1371/journal.pone.0090498

Shiell, M. M., Champoux, F., & Zatorre, R. J. (2015). Reorganization of Auditory Cortex in Early-deaf People: Functional Connectivity and Relationship to Hearing Aid Use. Journal of Cognitive Neuroscience, 27(1), 150–163. doi:10.1162/jocn_a_00683

Simon, M., Lazzouni, L., Campbell, E., Delcenserie, A., Muise-Hennessey, A., Newman, A. J., Lepore, F. (2020). Enhancement of visual biological motion recognition in early-deaf adults: Functional and behavioral correlates. PLOS ONE, 15(8), e0236800.doi:10.1371/journal.pone.0236800

Stephan, K. E., Penny, W. D., Moran, R. J., den Ouden, H. E. M., Daunizeau, J., & Friston, K. J. (2010). Ten simple rules for dynamic causal modeling. NeuroImage, 49(4), 3099–3109.doi:10.1016/j.neuroimage.2009.11.015

Stivalet, P., Moreno, Y., Richard, J., Barraud, P.-A., & Raphel, C. (1998). Differences in visual search tasks between congenitally deaf and normally hearing adults. Cognitive Brain Research, 6(3), 227–232.doi:10.1016/s0926-6410(97)00026-8

Striem-Amit E, Almeida J, Belledonne M, Chen Q, Fang Y, Han Z, Caramazza A, Bi Y. (2016) Topographical functional connectivity patterns exist in the congenitally, prelingually deaf. Sci Rep. 18;6:29375. doi: 10.1038/srep29375.

Vachon, P., Voss, P., Lassonde, M., Leroux, J.-M., Mensour, B., Beaudoin, G., Lepore, F. (2013). Reorganization of the auditory, visual and multimodal areas in early deaf individuals. Neuroscience, 245, 50–60. doi:10.1016/j.neuroscience.2013.04.004

Voss, P., Tabry, V., & Zatorre, R. J. (2015). Trade-off in the sound localization abilities of early blind individuals between the horizontal and vertical planes. Journal of Neuroscience, 35(15), 6051–6056.

Yamauchi, K., and Yamadori, T. (1982). Retinal projection to the inferior colliculus in the rat. Acta Anat (Basel). 1982;114(4):355–60. doi: 10.1159/000145608

